# The PGC-1α Axis Links Estrogen Receptor-α Signaling to Japanese Encephalitis Virus Replication in Neuronal Cells

**DOI:** 10.64898/2026.10.12.758457

**Authors:** Kiran Singh, Vishnu Kumar, Marek Bartkuhn, Sachin Kumar

**Affiliations:** Department of Biosciences and Bioengineering, Indian Institute of Technology Guwahati, Guwahati 781039, Assam, India; Biomedical Informatics and Systems Medicine, Science Unit for Basic and Clinical Medicine; Institute for lung health (ILH), Justus Liebig University Giessen

**Keywords:** Estrogen, Estrogen receptor alpha, Japanese encephalitis virus, *Flavivirus*, Neuroprotection, Neuroinflammation

## Abstract

Acute encephalitis due to the viral causative agents, such as Japanese Encephalitis Virus (JEV), remains a leading cause of viral encephalitis in Asia, contributing to high mortality and long-term neurological sequelae. The host factors that modulate JEV replication are not well understood. In this study, we investigated the modulation of JEV infection by estrogen *in vitro* and *in vivo*. The results demonstrated that estrogen enhances JEV replication in neuronal cells via estrogen receptor alpha (ERα) by modulating peroxisome proliferator-activated receptor gamma coactivator-1 alpha (PGC-1α). Since JEV pathogenesis increases cellular production demands, it results in dysregulation of cellular homeostasis, whereas estrogen treatment helps keep cellular health in check. Also, the mouse model exhibited an increased viral burden with protective antibody titers in response to estrogen. Conclusively, this work highlights the mechanistic link of estrogen-driven mechanisms and identifies the ERα-PGC-1α-axis as a potential target for sex-specific therapeutic interventions.

## Introduction

Acute encephalitis is a serious neurological condition marked by inflammation of the brain parenchyma. It leads to significant morbidity, mortality, and long-term neurological deficits. This includes encephalitis, acute encephalitis syndrome (AES), and autoimmune encephalitis (AIE). AES is a serious public concern, causing altered mental status with supporting signs like fever, seizures, CSF pleocytosis, or neuroimaging changes [1, 2]. Symptoms range from mild fever and headache to severe coma and death. These conditions arise from the diffusion of pathogens or autoantibodies in brain tissue, causing reversible or irreversible damage [3]. Despite advances, over half of cases remain unexplained, highlighting the need for improved diagnostic tools and surveillance strategies.

Among other etiological agents, viral encephalitis is largely caused by the Japanese encephalitis virus (JEV), causing neuroinflammation and long-term neurological impairment [4]. Japanese encephalitis (JE) is part of a “pure neurologic illness” spectrum within AES, distinct from multisystem syndromes caused by other agents. JE is associated with extrapyramidal signs and abnormal movements, classically seen in patients with Parkinson’s disease (PD) [5, 6]. The 30-50% symptomatic neuroencephalitis cases caused by JEV infections lead to permanent lifelong complications due to partial remission. Among those, 30% suffer significant fatality [7]. In a few cases, JEV causes bulbar paralysis due to injured cranial nerves, linking the symptoms to Guillain-Barré syndrome (GBS) [8].

JEV is a positive-sense RNA virus, part of the *Flaviviridae* family. Flaviviruses are known to have high energy demands [9], leading to an altered host response by subverting mitochondrial energy kinetics [10, 11], metabolic reprogramming [12–15], and immune evasion [16, 17]. JEV modulates neuronal death by caspase 3-mediated apoptosis, leading to a reduction in mitochondrial membrane potential [18, 19] and PTEN-Induced Kinase 1 (PINK1)-induced mitophagy [20]. Furthermore, JEV is associated with increased glycolysis, leading to catabolic patterns in lipid metabolism [21–23]. Also, an increase in ER stress due to the burden of unfolded protein response is concurrent with JEV infection [24]. The pathobiology of JEV leads to neuronal damage through virus-induced cellular death and to an inflammatory response by resident immune cells in the CNS [7].

Epidemiological trends of JEV infection patterns revealed consistent sex-based differences. Surveillance data from Taiwan (2010-2022) reveal that males account for 64% of JEV cases compared to 36% in females [25]. This male predominance is consistently observed across multiple Asian populations, with mainland China reporting a male-to-female ratio of 1.53:1 [26]. Although the male predominance doesn’t account for the higher prevalence of the disease in males, it includes several socio-economic reasons, such as the number of patients reporting, fewer female workers in the paddy fields in Asian countries, etc. Therefore, no experimental evidence is present to claim the actual sex based mechanism involved in these epidemiological trends.

Interestingly, the CNS region is protected from cellular abnormalities through crosstalk within the hormonal milieu, thereby maintaining structural and functional integrity. Estrogen plays an important role in brain activity mediated by two classical estrogen receptor α, β (ERs) and GPR30. The function of ERs is important for neuronal survival, metabolism, synaptic plasticity, and the modulation of memory-related gene expression, which are mediated largely through both genomic and non-genomic pathways [27, 28]. Moreover, ERs functioning depends on the synchronized activity of estrogen, coregulators, and post-translational modifications (PTMs) [29]. Estrogen activates MAPK/ERK and Akt pathways to improve cell proliferation, survival, and energy metabolism. It indirectly controls the mitochondrial function in neurons through the ATP production, biogenesis, and transcription of mitochondrial genes [30, 31]. The cooperation exists between estrogen and Peroxisome proliferator-activated receptor γ coactivator 1α (PGC-1α) for managing mitochondrial function to promote energy metabolism [32]. It is a key coactivator that controls mitochondrial biogenesis, metabolic regulation, and cellular differentiation [33, 34].

Estrogen and ERα are involved in the neuroprotective and neurotoxic functions, which depend on several factors [35, 36]. Estrogen also supports the innate immune responses; it helps to dampen excessive NF-κB activation [37] while supporting type I interferon production [38, 39]. Dysregulation of ERα signaling contributes to various pathological conditions, including endocrine-resistant breast cancer [40], and neurodegenerative diseases such as Alzheimer’s (AD) [41, 42]. Women are also shown to have a lower risk of PD, but can develop chronic symptoms faster than men [41, 43–45]. Thus, ERα acts as a critical molecular integrator of estrogenic signaling in both health and disease.

Few reports support the role of sex hormones in viral diseases such as ZIKA, Hepatitis E, and HCV [14, 38, 46–49]. The severity of any disease and the persistence of symptoms, as in the case of JEV, depend on the interplay within core cellular metabolism. Estrogenic control of nuclear and mitochondrial genes, and the molecular pathways that govern organelle dynamics, is crucial for cellular maintenance [30, 31]. Despite extensive studies on JEV-induced neuroinflammation, the role of estrogen and ERα is largely unknown in this context. This study has important clinical implications as therapeutic options for JE are limited. So, elucidating the role of estrogen signaling in JEV infection may therefore provide insights into therapeutic strategies and antiviral interventions.

Based on the gap in understanding, we examined the hypothesis that estrogen and its signaling through ERα affect JEV growth kinetics. The study investigated transcriptome reanalysis, overexpression, and downregulation, along with key modulators in JEV-infected neuronal cells. The response was also studied at the organism level in the mouse model. The data obtained in the study suggested that estrogen and its associated pathway are key modulators of JEV infection.

## Materials and Methods

### In silico analysis

RNA-sequencing data associated with the Gene Expression Omnibus (GEO) accession GSE221680 of JEV-infected SH-SY5Y neuronal cells were downloaded from the NCBI database for downstream analysis. The nf-core RNA-seq pipeline (v3.14.10) was implemented in NEXTFLOW (v24.04.2) with default parameters in Docker mode for quality control, filtering, trimming, and gene-specific count table generation. Reads were aligned with the Homo sapiens GRCh38 genome and gene annotations from Illumina’s iGenome repository. One control sample (Control_2) containing only single-end reads was excluded from downstream analyses to maintain consistency across the dataset. Lastly, we used the 2 control (Control_1 and Control_3) and 3 infected (Infected_1, Infected_2, and Infected_3) samples for downstream analysis.

Differential gene expression analysis was performed in R using DESeq2. For volcano plot generation, the EnhancedVolcano R package was used. Differentially expressed genes were defined using a p-value < 0.05 and an absolute log2 fold change threshold of > 1. For heatmap visualization, differentially expressed genes were ranked by adjusted p-value, and the top 100 genes were selected for plotting using the ComplexHeatmap R package. Pathway enrichment analysis was performed using the clusterProfiler package in R. Gene Set Enrichment Analysis (GSEA) was conducted using genes ranked according to stat derived from DESeq2 results to identify enriched pathways against MSigDB hallmarks and Reactome databases. Pathways with a normalized enrichment score (NES) and p-value < 0.05 were considered significantly enriched, with positive and negative NES values indicating upregulation and downregulation, respectively. Gene-pathway networks were constructed from Reactome GSEA results and visualized in R using ggraph, ggplot2, ggnewscale, and dplyr. Networks were arranged using the Kamada-Kawai layout.

### *In vitro* analysis Cells and Virus

The mouse neuronal cells (N2a) and baby hamster kidney cells (BHK-21) were procured from the National Centre for Cell Science, Pune, India. All the cell lines were routinely maintained in appropriate medium; N2a and BHK-21 were maintained in Dulbecco’s modified eagle medium (DMEM), containing 1X penicillin-streptomycin antibiotic with 10% fetal bovine serum (FBS) in 5% CO2 at 37°C. In experiments involving treatment with various estrogen receptor-modulating ligands, cells were treated in phenol red-free DMEM-F12 supplemented with penicillin-streptomycin and 10% csFBS (Charcoal-stripped Fetal Bovine serum).

The JEV strain SA_14_-14-2 (GenBank accession number JN604986) belongs to genotype III and was obtained from the Department of Health and Family Welfare, Government of Assam, India. The JEV stock preparation and titration were performed in BHK-21 cells. BHK-21 cells at 80% confluency were infected with JEV for stock preparation. JEV titers were calculated by TCID_50_ and by plaque assays following the standard protocol. The MOI was calculated using the standard infective virus particles-to-cell ratio, where cells were counted by an automatic cell counter.

### Plasmids, antibodies, and reagents

The plasmids were used for ERα-related studies: pEGFP-C1-ER alpha (Addgene #28230) and 3xERRE/ERE-luciferase (Addgene #37852). The anti-JEV-NS1 (GTX633820, GeneTex, USA), anti-JEV-E (GTX125867, GeneTex, USA), anti-ERα (sc-8005, Santa Cruz, USA), anti-PGC-1α (A11971, Abclonal, USA), anti-GAPDH (97166, Cell Signaling Technology, USA), anti-βactin (4967, Cell Signaling Technology, USA), Alexa Fluor^TM^ 594 goat anti-rabbit IgG (H+L) (2018240, Invitrogen), Alexa Fluor^TM^ 594 goat anti-mouse IgG (H+L) (1826426, Invitrogen), goat anti-rabbit (Invitrogen, USA), and anti-mouse (Cell Signaling Technology, USA) conjugated with HRP antibodies were used for various cell based studies to see the levels of protein expression.

The 17β-estradiol (E8875, Sigma-Aldrich, USA), Tamoxifen (sc-3542C, Santa Cruz, USA), Fulvestrant (sc-203435, Santa Cruz, USA), 2-Methoxy-estradiol (454180, Merck, USA and Canada), Charcoal-stripped serum Fetal Bovine Serum (F6765, Sigma-Aldrich, USA) reagents were used for the experiments related to cells.

### Cell viability and virus growth kinetics

N2a cells were seeded at a density of 2 × 10^6^ cells/well in a six-well plate and incubated for 12 h at 37°C, 5% CO_2_ incubator. Once the cells were attached, the media was changed with Phenol-red-free DMEM-F12 supplemented with 2% of csFBS, subsequently JEV infection was given at 24 h. The cell viability in control and infected cells was assessed using the Trypan blue assay. Temporal virus growth kinetics were calculated at 0.01, 0.1, and 1 MOI by using the TCID_50_ assay.

### Cell treatment and analysis

The N2a cells were treated with different concentrations of Estrogens at 18 h (0.5 nM, 2 nM, 10 nM, 20 nM, and 40 nM), followed by 0.1 MOI JEV infection at 24 h. Similarly, treatment of other ER alpha modulators was done similarly at appropriate concentrations. Tamoxifen (Tam), Fulvestrant (Ful), and 2-methoxyestradiol (2-MeOE2) were used at 1 μM, 200 nM, and 50 nM concentrations, respectively.

For overexpression experiments, pEGFP-C1-ER alpha was transfected using Lipofectamine 3000 (Invitrogen, USA) according to the manufacturer’s instructions. Similarly, the nuclear cytoplasmic fractionation was performed according to the manual instructions. Briefly, the cells were collected in PBS, and the pellet was resuspended in CE buffer. The supernatant containing the cytoplasmic extract was collected after 3 minutes of incubation on ice and centrifugation. The pelleted nuclei were washed with CE buffer without detergent and resuspended in NE buffer, and glycerol was added to the CE tube to 20%. The collected fractions were then stored at -80 °C.

For luciferase activity, the pEGFP-C1-ER alpha and 3xERRE/ERE-luciferase were transfected in a 3:1 ratio, and the luciferase activity was measured using Thermo Scientific Varioskan LUX.

### Gene, protein expression, and virus infectivity analysis

Total RNA was isolated using RNAiso reagent (Takara, Japan) and converted into cDNA using iScript cDNA Synthesis Kit (Biorad, USA). Quantitative real-time PCR analysis was carried out in the QuantStudio5 system (Applied Biosystems, USA) using a Power-up SYBR master mix (Invitrogen, USA). GAPDH was used as an internal control, and values are represented as mean fold change with respect to the cell control (Mock). Protein estimations were carried out using western blotting, keeping GAPDH, β-actin, and H3 histone as a loading control for the experiments. The relative quantification was done using ImageJ software. For infectivity quantification, supernatants from the treated samples were serially diluted in DMEM without FBS. Virus adsorption was performed by taking 100 µL of diluted supernatant for 2 h. After adsorption, cells were incubated for 72 h in 0.8% methylcellulose DMEM supplemented with 2% FBS. Cells were stained with crystal violet for visualization. Virus quantification is done at an appropriate dilution, and representative images are shown.

### *In vivo* analysis

A total of 3-4 weeks old 12 Swiss Albino female mice were randomly allocated into four experimental groups of three animals each (n = 3 per group). Control group animals receiving no estrogen and no virus; estrogen group animals receiving estradiol only; JEV group animals receiving intracerebral viral inoculation only; and (iv) estrogen and JEV (Estrogenvirus) animals receiving both estradiol and intracerebral viral inoculation. Estradiol was administered subcutaneously (2.5 µg per animal) as a single priming dose on day 0, dissolved in sterile sesame oil. Control and JEV only groups received an equivalent volume of vehicle by the same route and schedule. JEV was administered by intracerebral injection on day 1 after 12 h of estrogen treatment, designated as day 0. A 10^6^ pfu/ml viral titer was delivered. Control and estrogen-only groups received an equivalent volume of sterile PBS by the same route. Blood was collected on days 0, 3, 5, and 14 via retro-orbital under anesthesia. At each time point, blood was divided into two aliquots: one was collected into EDTA-coated vacutainers and centrifuged to isolate plasma; the second aliquot was collected into serum-separator tubes, allowed to clot at room temperature, and centrifuged under the same conditions to obtain serum. Aliquoted plasma and serum were stored at -80 °C until analysis. The animals were euthanized by cervical dislocation following terminal bleeding. The brain and spleen were rapidly excised. Brain tissue was processed for histological analysis (H&E staining and IF-IHC), while spleen tissue was snap-frozen for downstream protein analysis. Tissues were stored at -80 °C until processing.

### Clinical observations and body weight

Over the 14-day observation period, survival and visible signs such as ruffled fur/hunched posture were examined. Body weight was recorded at days 0, 3, 5, and 14.

### Serum neutralization test (SNT)

Virus-neutralizing activity in serum was assessed using a varying serum concentration for a constant virus neutralization test on BHK-21 cells in a 96-well plate. Serum samples were heat-inactivated at 56 °C for 30 min and serially two-fold diluted, starting from 1:10. Equal volumes of diluted serum and working virus dose 100 TCID50 of the virus were mixed and incubated at 37 °C for 2 h, then added to sub-confluent monolayers. After 3 to 5 days at 37 °C with 5 % CO2, infection was scored based on cytopathic effect. Neutralizing titers were expressed as the reciprocal of the highest serum dilution yielding ≥ 50 % reduction in CPE. Each sample was assayed in triplicate.

### Histology analysis

#### Hematoxylin and eosin (H&E) staining

Brain tissue was fixed in 4 % paraformaldehyde in PBS for 24h at 4 °C, dehydrated through a graded ethanol series, cleared in xylene, and embedded in paraffin. Sections were cut on a microtome and mounted on poly-L-lysine-coated slides. Sections were deparaffinized, rehydrated, and stained with hematoxylin and eosin according to standard protocols. Slides were examined under an upright Olympus Fluorescence microscope (CKX53).

#### Immunofluorescence and immunohistochemistry (IF-IHC)

Paraffin-embedded brain sections were deparaffinized and subjected to heat-induced antigen retrieval in citrate buffer pH 6.0 for 20 mins. Sections were permeabilized in 0.1 % Triton X-100 in PBS and blocked with 5 % goat serum in PBS-T for 1 h at room temperature. A cocktail of Primary antibodies against JEV E and NS1 was applied overnight at 4 °C. After washing, sections were incubated with fluorophore-conjugated secondary antibody. For immunofluorescence, nuclei were counterstained with DAPI. Images were acquired on an EVOS Floid by Thermo Fischer Scientific using identical acquisition settings across groups. Quantification was performed in the software ImageJ.

### Statistical analysis

The data presented in the results are the average of three independent biological experiments. Group comparisons were performed by one-way ANOVA with Tukey’s post hoc test for normally distributed data and two-way ANOVA with Tukey’s multiple comparisons test on GraphPad Prism 8. Data are expressed as mean±SD, and significance is represented as *P < .05, **P < .01, ****P < .0001, n.s., not significant.

## Results

### *In silico* transcriptomic analysis

To understand the molecular mechanism associated with JEV infection, the publicly available transcriptomic dataset (GSE221680) of neuronal cells corresponding to 48 h post-infection was analyzed. Differential expression between infected and control neuronal cells reflected 465 up-regulated and 118 down-regulated genes among 29744 analyzed genes (Figure 1A). Among the top 100 differentially expressed genes, highly upregulated genes were associated with antiviral (e.g., NFKB2, ISG15, DDX58) and ER-stress (e.g., HSPA5, DDIT3) responses. On the other side, DNA modification and repair (e.g., UHRF1) and cell cycle (e.g., CCND1) regulated genes were downregulated (Figure 1B). Hierarchical clustering of the top differentially expressed genes (DEGs) (Figure 1B) resolved an up-regulated antiviral, ER-stress, and cell proliferation response. GSEA of MsigDB Hallmark confirmed this pattern at the pathway level; Interferon-γ/α response, TNFα-NF-κB signaling, and the unfolded protein response were the top positively enriched sets, while E2F targets, G2M checkpoint, MYC targets, and mitotic spindle were among the most negatively enriched gene sets (Figure 1C). In addition, the Hallmark Estrogen response early gene set was significantly associated with downregulation of gene expression in infected cells.

**Figure 1:**
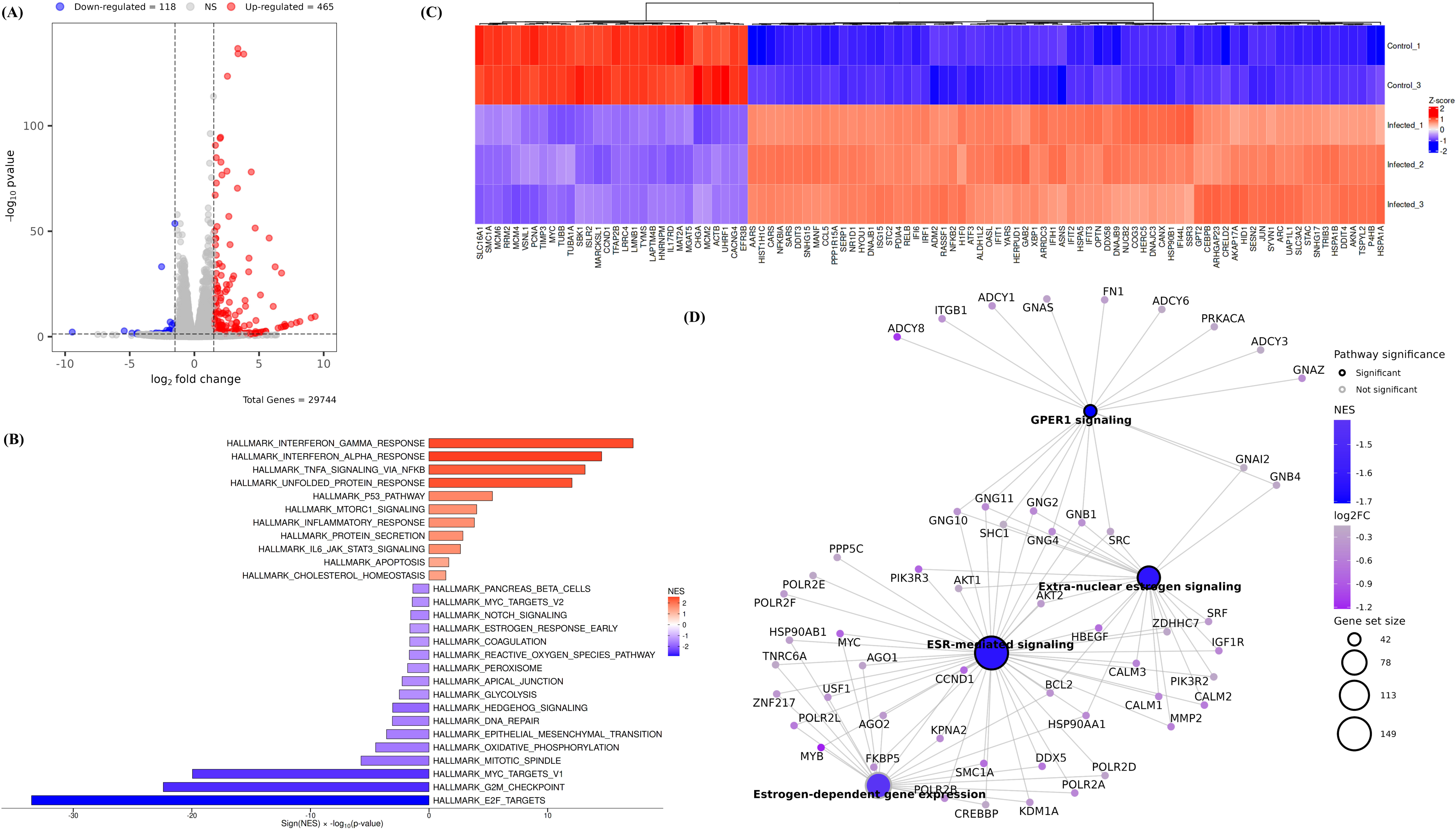
Transcriptomic analysis reveals estrogen-mediated gene expression networks. Volcano plot showing differentially expressed genes (DEGs) between control and JEV-infected neuronal cells **(A)**. Significantly upregulated genes (log₂FC > 1, padj < 0.05), downregulated genes (log₂FC < -1, padj < 0.05), and non-significant changes are represented as red, blue, and gray dots, respectively. Heatmap of the top 100 most significantly modulated genes in infected neuronal cells compared to control, with hierarchical clustering **(B)**. The color scale represents z-score normalized expression values. MSigDB Hallmark Gene Set Enrichment Analysis (GSEA) showing significantly modulated biological processes upon JEV infection in neurons **(C)**. Bar plot displays various hallmark gene sets with enrichment scores, highlighting interferon responses (both alpha and gamma), inflammatory pathways, and metabolic processes as prominently affected pathways. Gene-pathway network showing the associations between genes and enriched pathways. Pathway node size represents gene-set size, pathway color represents NES, gene-node color represents log2 fold change, and pathway border color indicates adjusted P-value significance (≤0.05) **(D)**.

Further analysis using Reactome-based GSEA, visualized through a gene–pathway network plot, revealed significant negative enrichment of pathways associated with extranuclear estrogen signaling, GPER1 signaling, and ESR-mediated signaling (Figure 1D). Although the estrogen-dependent gene expression pathway did not reach statistical significance, its leading-edge genes showed substantial overlap with those contributing to the significantly negatively enriched ESR-mediated signaling pathway. These overlapping genes included the canonical ERα transcriptional targets CCND1 and MYC, key regulators of cell-cycle progression and proliferation, as well as several ERα coregulators and chaperones, including HSP90AA1, DDX5, CREBBP, KDM1A, and FKBP5.

### Effect of estrogen on neuronal cells

Our transcriptomic analysis revealed downregulation of estrogen signaling-related genes following JEV infection, prompting us to investigate whether estrogen (E2), a ligand of estrogen receptors, modulates JEV replication. To address this, neuronal cells were pretreated with different concentrations of E2 (0.5-40 nM), which showed upregulation of JEV replication, with significant increases in mRNA levels of ∼6-, 8-, and 15-fold at 10, 20, and 40 nM, respectively. Similarly, viral protein showed upregulation of ∼1.7-, 3.7-, and 4.6-fold at 10, 20, and 40 nM, respectively (Figure 2A and B). JEV protein and titration also established these findings, showing a ∼5.5-fold and ∼7.5-fold increase at a 40 nM concentration of NS1 protein and infectious viral particles, correlating with the E2 dose (Figure 2C-D).

**Figure 2:**
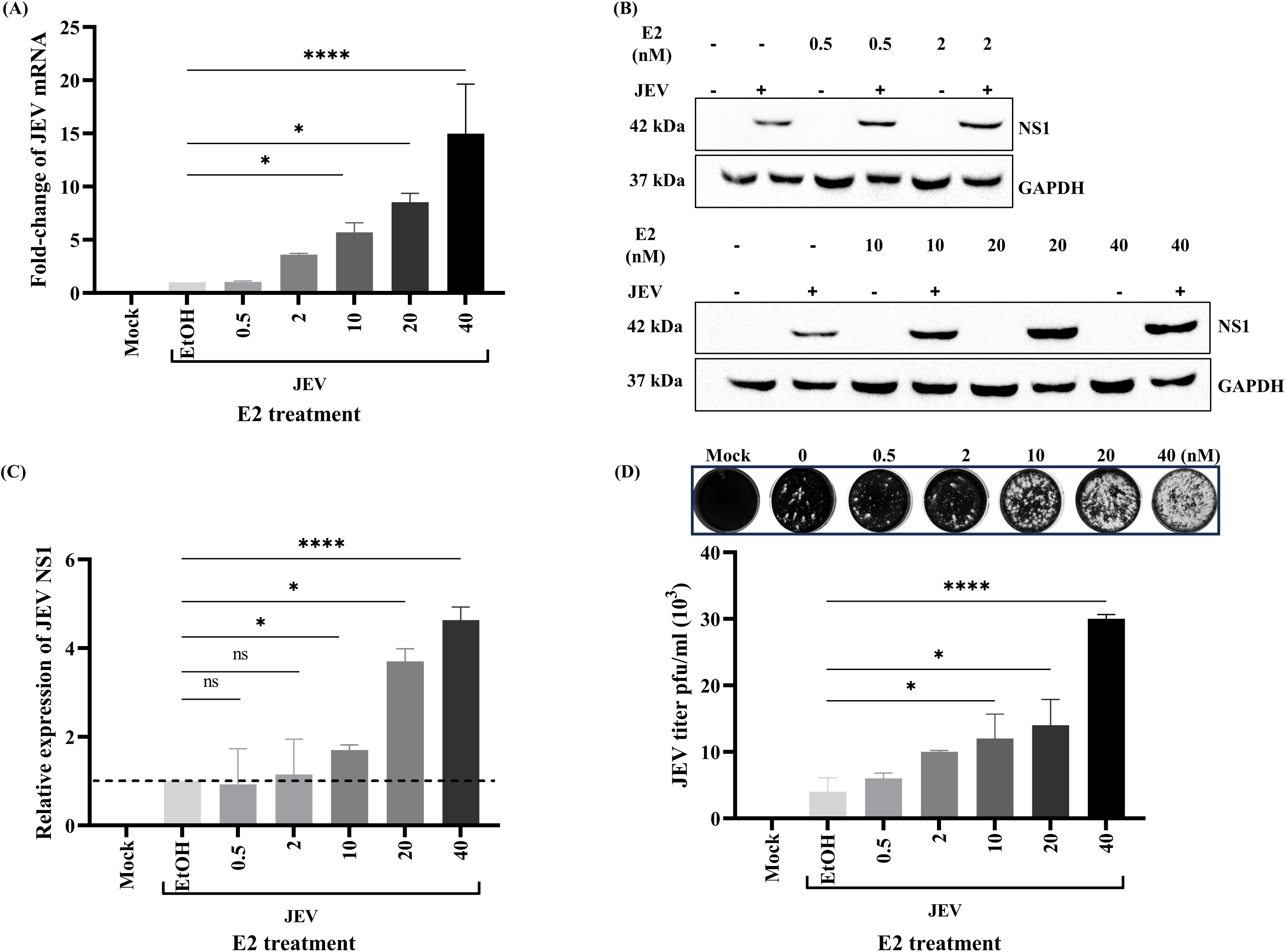
Estrogen dose-dependent enhancement of JEV replication in N2a cells. qRT-PCR quantification showing fold-change (fold change w.r.t JEV control) in JEV mRNA expression at different E2 concentrations **(A).** Western blot analysis of JEV protein and GAPDH expression across E2 doses **(B).** Densitometric quantification (fold change w.r.t JEV control) of JEV protein bands from the panel, normalized to GAPDH loading control, confirming dose-dependent viral protein concentration with statistical significance achieved at concentrations ≥10 nM E2 **(C)**. The JEV-only control was present on both blots and used to normalize densitometry across membranes. Representative plaque assay images are shown, and quantification of viral titers (pfu/mL) was done at an appropriate dilution. The absolute JEV titer determined by plaque assay **(D)**. Data represent mean±SD from three independent experiments.

### Role of ER alpha in JEV infection

To investigate the role of ER alpha in JEV infection, N2a cells were infected with JEV and analyzed after 72 h. JEV infection was first confirmed at the mRNA level by qPCR (Figure 3A). Infected cells showed an approximately 95% reduction in ERα mRNA expression compared with control cells (Figure 3B). Similarly, a 50% reduction in ERα protein levels was evident upon JEV infection (Figure 3C-D). Further, N2a infected with different MOI of JEV showed a gradual increase of ERα mRNA (∼1.75) up to 48 h (2-fold) and then a decrease at 72 h in both 0.1 (∼1.8 fold) and 1 (∼1.5 fold) MOI (Figure 3E). At 0.01 MOI (marked by black dashed lines, Figure 3E), there are no significant changes in the ERα mRNA. The increase in virus titer through 24 h, 48 h, and 72 h is consistent with MOI (Figure 3F). A similar pattern of ERα as with mRNA was observed in protein levels as well, where it showed an increase in 48 h (∼5.7-fold) and then a decrease at 72 h (up to ∼ 3.5-fold) (Figure 3G-H). Subcellular fractionation of mock and infected neuronal cells demonstrated that JEV infection led to cytoplasmic depletion of ERα, in parallel with enhanced cytoplasmic accumulation of viral proteins (Figure 3I and 3J).

**Figure 3:**
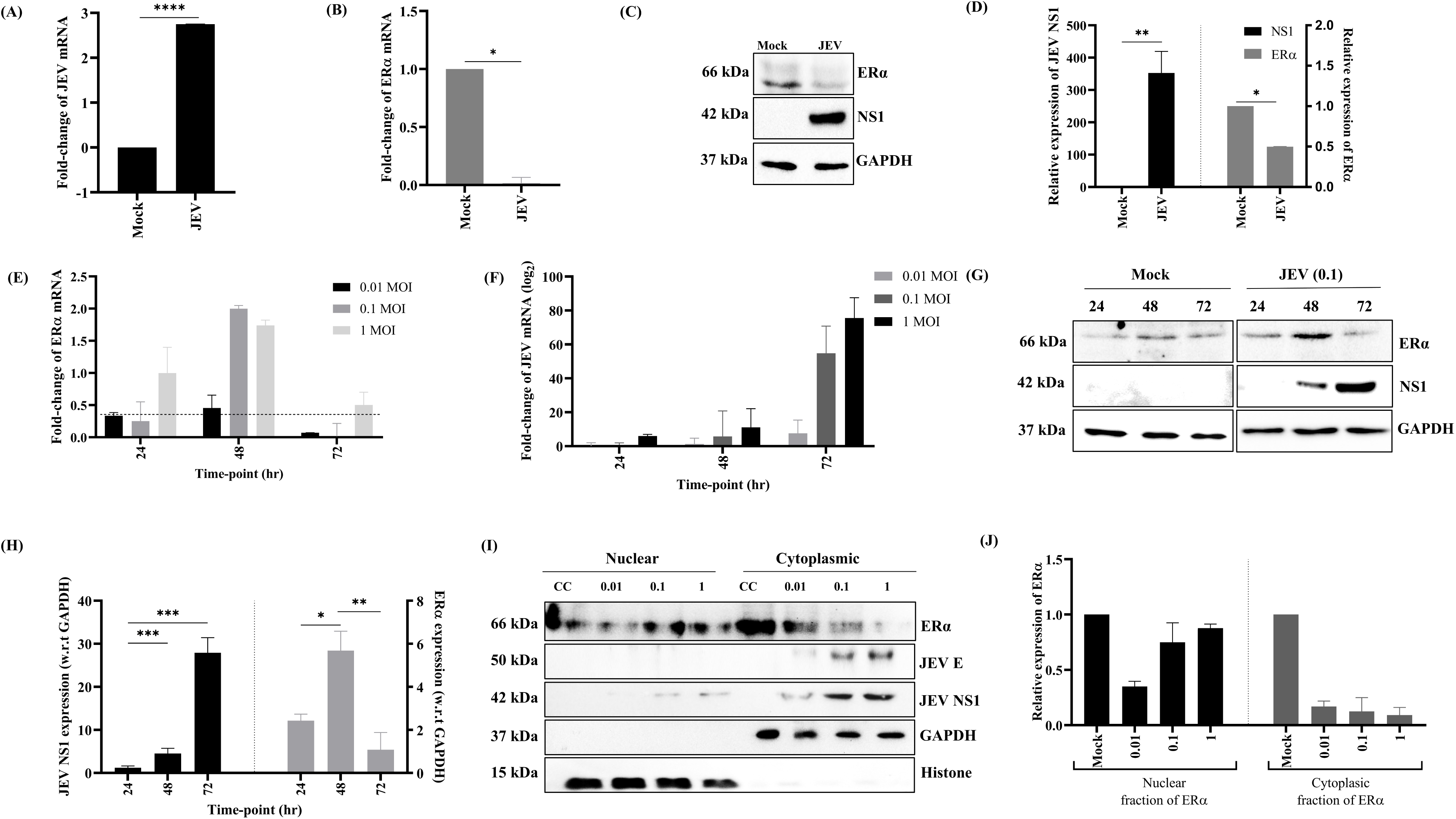
JEV infection modulates estrogen receptor α expression in a time and MOI-dependent manner. qRT-PCR showing mRNA of JEV (log10 fold change) and ERα upon JEV infection in N2a **(A-B).** Western blot and quantification of ERα protein and JEV protein expression **(C-D)**. Time- and MOI-dependent analysis of JEV replication and ERα expression **(E-F).** Western blot and quantification analysis (w.r.t. control) represent the same **(G-H)**. Western blots and their quantification of subcellular fractionation of ERα following infection **(I-J)**. Data represent mean±SD from three independent experiments.

### Effect of ERα modulation on JEV replication

N2a cells were further assessed for treatment with ERα antagonists tamoxifen (Tam) and fulvestrant (Ful) at 1 μM and 200 nM, respectively, prior to viral infection. Gene-expression data revealed that inhibition of ERα signaling significantly decreased JEV mRNA expression to ∼0.18-fold and ∼0.5-fold in Tam and Ful-treated cells, respectively, compared with controls (Figure 4A). Protein analysis confirmed a marked reduction in the JEV NS1 protein in both Tam and Ful-treated cells (Figure 4B and 4C). Plaque assays further showed reduced JEV titers by approximately 82% in Tam, and 45% in Ful-treated N2a cells (Figure 4D).

**Figure 4:**
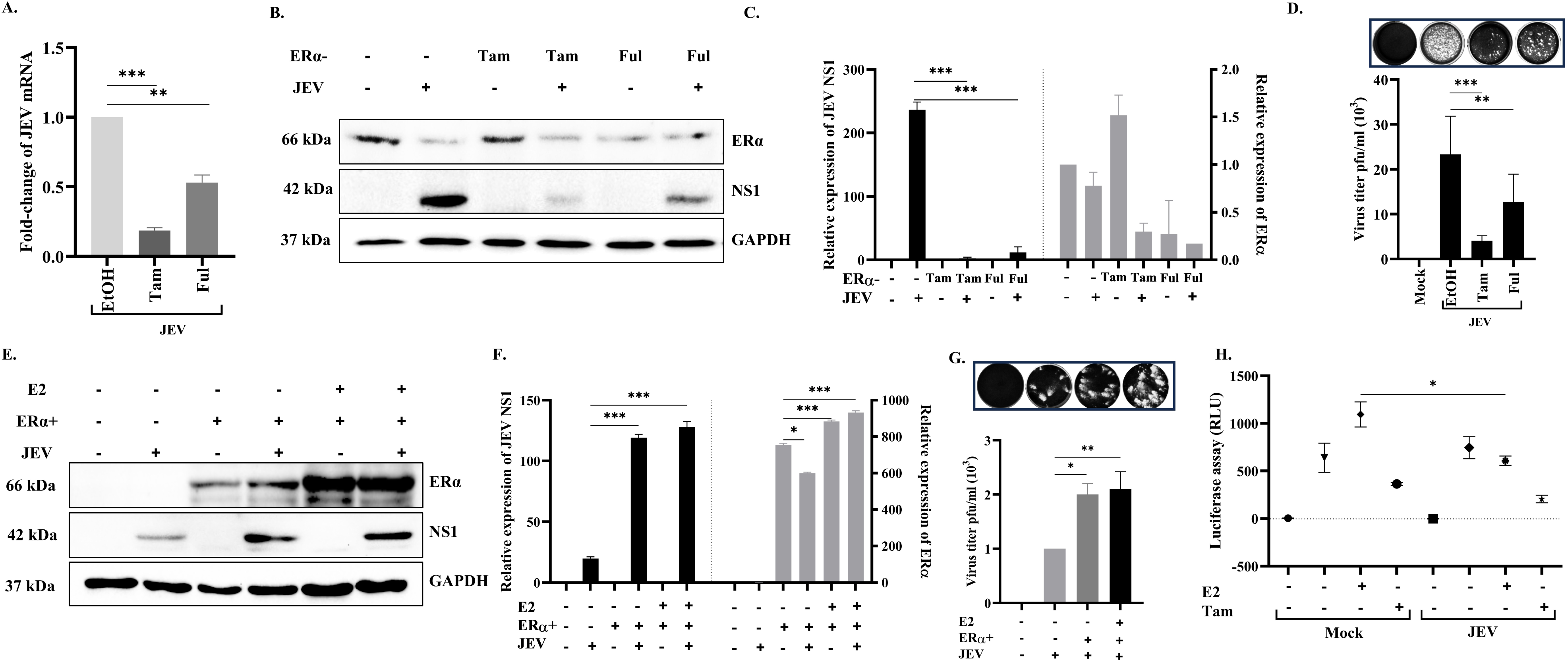
Estrogen receptor modulation affects JEV replication. qRT-PCR analysis showing relative fold-change in JEV mRNA expression (log2 fold change) following treatment with Tam or fulvestrant Ful prior to infection **(A)**. Western blot and its quantification analysis (compared with control) depicting ERα and JEV protein expression, with GAPDH as loading control **(B-C)**. Plaque assay quantification of viral titers (pfu/mL) with representative plaque images is shown above **(D)**. Western blot, quantification (compared with control), and plaque assay with viral titers quantified in the bar graph **(E-G)**. Luciferase assay upon overexpression of the ERα plasmid in N2a cells with controls **(H).** Data are expressed as mean±SD from three independent experiments.

To further confirm the role of ERα in JEV replication, the N2a cells were transfected with an ERα-overexpressing plasmid. The overexpression of ERα increased the JEV protein and viral titers. In addition, when estrogen supplementation was applied to ERα overexpressed cells, a subtle increase in the viral titers was observed compared to JEV-infected-transfected cells (Figure 4E, 4F, and 4G). Luciferase assay carried out with 3xERRE/ERE-luciferase and plasmid showed a significant decrease in the luciferase activity in JEV-infected cells supplemented with estrogen (Figure 4H). However, there is a decrease in infected cells, supplemented with Tam.

### Mechanistic insight into PGC-1α

To explore whether E2 influences cellular functions during JEV infection, such as mitochondrial biogenesis, the expression of peroxisome proliferator-activated receptor gamma coactivator-1 alpha (PGC-1α) was examined. Gene expression analysis showed that E2 treatment significantly upregulated (∼1.6-fold) PGC-1α mRNA compared to untreated control cells (Figure 5A). In contrast, JEV infection alone led to a substantial reduction (∼0.5-fold) in PGC-1α expression (Figure 5A). Interestingly, treatment with E2 and JEV partially restored PGC-1α expression (∼0.75-fold). Protein analysis demonstrated increased PGC-1α protein levels (∼5-fold) in E2-treated N2a cells, with a moderate recovery in E2 JEV-treated samples (∼4.2-fold) relative to mock-infected cells (Figure 5B-C). 2-MeOE2 treatment, a known PGC-1α inhibitor, showed ∼20% downregulation of JEV mRNA (Figure 5D), and significant decrease of PGC-1α mRNA in 2-MeOE2 treated (∼0.2 fold) and JEV infected cells (∼0.3 fold) (Figure 5E). Similarly, JEV protein (∼54%) and virus titer (∼52%) are reduced with treatment of 2-MeOE2 (Figure 5F-H).

**Figure 5:**
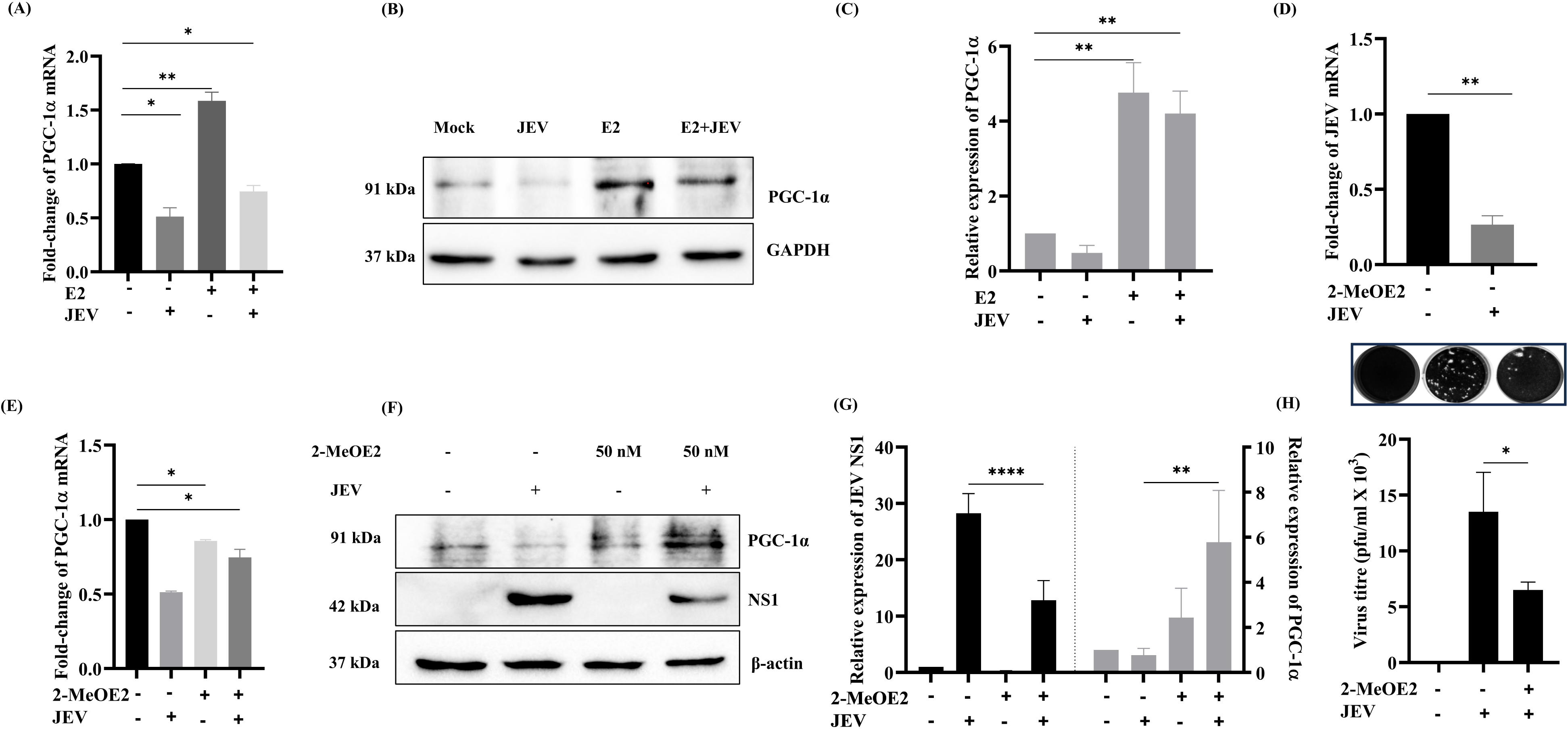
Estrogen enhances PGC-1α signaling during JEV infection. PGC-1α mRNA expression under E2 treatment and JEV-infected conditions **(A)**. Western blot and quantification analysis showing PGC-1α protein expression in N2a cells with controls **(B-C).** PGC-1α and JEV mRNA expression with treatment of 2-MeOE2 **(D-E)**. Western blots and quantification showing PGC-1α and JEV protein levels **(F-G)**. Plaque assay and quantification showing viral titers **(H)**. Data represent mean±SD from three independent experiments.

### E2 treatment studies in mice

The mice treated with E2 and infected with JEV did not show a drastic body weight change as compared to the control. Body weight was slightly decreased in the estrogen-virus and virus-only groups on the 5^th^ day. However, no clinical signs or mortality were observed in treated groups till day 14 (Figure 6A). Serum neutralizing titer (SNT) against the JEV was assessed on days 0, 3, 5, and 14. Serum samples collected from all groups on the 0^th^ Day showed a seronegative baseline. The estrogen-treated and JEV-infected mice group showed higher SNT on day 14. The control and estrogen-only groups remained seronegative at all time points (Figure 6B). Western blot analysis of spleen lysates showed that JEV E was upregulated in the estrogen-treated JEV group. (Figure 6C).

**Figure 6:**
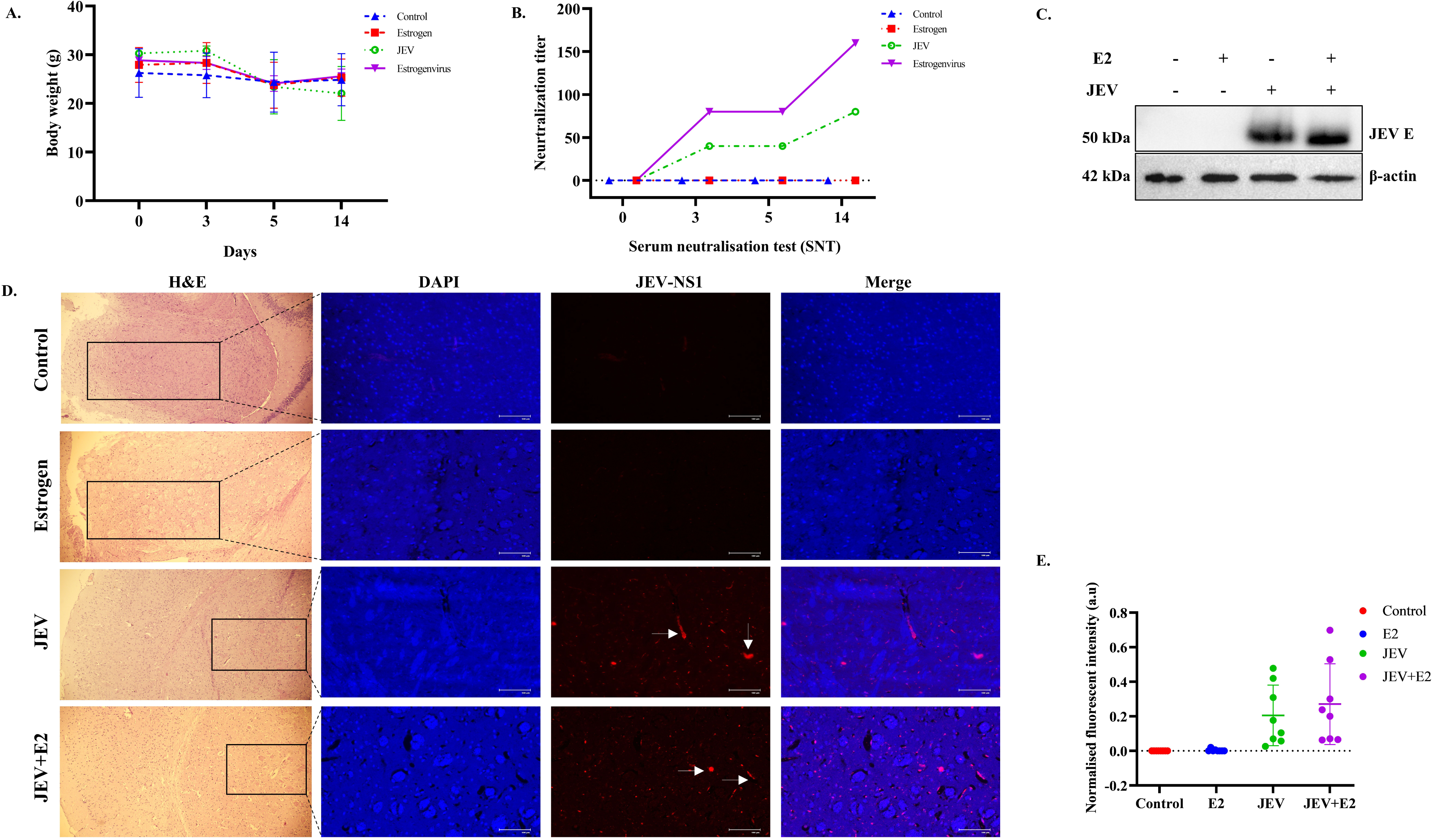
JEV virus effect in estrogen-treated mice. Body weight changes over time in Swiss-Albino mice following JEV infection, comparing control mice (blue line), estrogen-treated mice (red line), JEV-infected mice (green line), and estrogen-pretreated then JEV-infected mice (purple line) **(A)**. Serum neutralization test (SNT) across experimental groups at 0, 3, 5, and 14 days post-infection **(B)**. Western blot and quantification analysis of JEV envelope E protein expression in the spleen **(C)**. Quantification of normalized JEV NS1 fluorescence intensity (arbitrary units, a.u.) in brain tissue sections from JEV-infected (red) and JEV+E2-treated (blue) mice, derived from panel **(D).** Representative histopathological and immunofluorescence analysis of brain tissue sections from Control, Estrogen, JEV-infected, and JEV+E2-treated mice **(E)**. Each data point represents an individual field of view (n = 8 per group). Data are presented as mean±SD.

Histopathology of brain tissue sections revealed intact cytoarchitecture with no inflammatory infiltrate, and normal neuronal morphology was evident in the control group. Animals in the estrogen-only treatment group showed findings similar to those in the control group. In contrast, JEV-inoculated animals exhibited isolated areas of neuronal vacuolation and neutrophil infiltrates. Although the tissue architecture was intact, the estrogen-virus group showed greater vacuolation and infiltration than the virus-alone group. Immunostaining performed on mouse tissue sections showed more JEV antigen in the hippocampal region as compared to the control (Figure 6D-E).

## Discussion

AES cases predominantly dominate with JEV-related pathophysiologies [50]. JEV has developed an effective mechanism to invade and suppress the host immune system to establish the infection in the CNS [51, 52]. Our understanding of JEV pathogenicity has increased in recent years; however, the underlying molecular mechanism regarding its neurotropism remains unclear. Host gene transcription plays a vital role in cellular homeostasis. The transcriptomic analysis of human neurons infected with JEV showed upregulation of genes involved in inflammation, ER stress, and apoptosis. In contrast, genes involved in DNA replication and the cell cycle were downregulated during viral replication [53, 54]. This is an expected outcome in an infected cell to tune towards antiviral defense rather than cell division. The basic understanding of modulation in signaling pathways can be used to pinpoint major targets for anti-viral strategies.

Our gene enrichment analysis showed downregulation of estrogen-signaling genes in neurons upon JEV infection. ERα functions as a ligand-activated transcription factor that recruits coregulators and chaperones to estrogen-response elements, thereby controlling target genes governing proliferation, survival, and metabolism [55]. The availability of the coregulators reshapes the estrogen-responsive transcriptional output. Our data supported the involvement of multiple ER-targeted genes, cumulatively deregulating the estrogen signaling. It could affect major cellular metabolic pathways if dysregulated. We hypothesize that JEV-infected neuronal cells may exhibit dysregulated estrogen signaling and estrogen-response signaling.

The replication kinetics of JEV depend on multiple factors, including tissue tropism, viral load, and underlying metabolic disorders. Our data suggested that estrogen has a positive effect on JEV replication. The earlier work reported that pregnant women and fetuses are more susceptible to the Zika virus and SARS-CoV-2 infections [56, 57]. In contrast, estrogen suppresses HIV latency and influenza infections [58, 59]. Nevertheless, E2 has multiple functions in regulating reproduction, homeostasis, neurotransmitter signaling pathways, neural growth, and synaptic responses [60–64]. Moreover, ERα has been shown to contribute to cancer progression [65, 66]; however, its role in viral infection remains poorly characterized. We hypothesized that genes involved in ERα signaling in the mouse neuroblastoma cell line N2a might have implications for JEV biology. Presumably, modulation of cellular pathways via ERs can have a visible effect on neuroinflammation, as shown earlier [65].

The ERα signaling is generally regulated through estrogen [55]. The cumulative effect that we observe on JEV replication due to E2 treatment could pinpoint the involvement of ERα in neuronal cells. Our investigation of ERα modulation in JEV-infected neuronal cells over the time course confirmed that its levels declined as the viral infection progressed. Our data also suggested upregulation of ERα during the early phase, which reiterates its neuroprotective function. Furthermore, infection at higher MOI suggests that increased viral loads significantly affect ERα. This modulation suggests that JEV manipulates estrogen signaling, potentially by sequestering ERα or its transcriptional activity at higher titers. The data indicate that ERα is neuroprotective during the early phase of viral replication, whereas its levels are reduced at later stages. Perhaps JEV infection might be channelizing the ERα towards its own benefit. ERα sequestration has been shown to be beneficial in Alzheimer’s disease, whereas GPR30 mitigates oxidative stress in neurological diseases [66, 67]. Moreover, lower ERα levels could reduce cell sensitivity to E2 by reducing the number of available receptors for signaling [68]. This desensitization reduces E2-driven protective effects and antioxidant gene programs, leaving neurons more susceptible to oxidative stress and injury [69]. Nuclear ERα translocation regulates transcription of ERE-driven genes and promotes mitochondrial biogenesis, antioxidant defense, and cellular resilience [70]. Presumably, sequestration of ERα in the cytoplasm upon JEV-infection would suppress these protective pathways, diminish neuronal energy metabolism, and may divert resources toward its own replication.

Pharmacological inhibition of ERα signaling markedly reduced JEV replication, suggesting that ERα is a critical modulator of JEV replication. Tam binds to ERα, which can inhibit transcriptional activation, whereas Ful binding leads to ubiquitin-mediated degradation of ERα [71]. The stronger antiviral response to tam treatment suggests that interference with ERα signaling, rather than receptor degradation alone, exerts a more pronounced inhibitory effect on JEV replication. These results collectively indicate that ERα and its downstream signaling pathways contribute to JEV replication. Similar biology was reported for the Zika virus, in which Zika exploits estrogen-driven signaling, which was reduced by Tam’s modulatory role on ERα [72]. Furthermore, transient ERα overexpression enhances viral replication, suggesting that ERα either directly or indirectly facilitates a cellular environment conducive to JEV replication, perhaps by altering cellular homeostasis.

JEV replication in neuronal cells leads to calcium dysregulation, suggesting the accumulation of damaged mitochondria and subsequent impairment [20, 73]. DENV and ZIKV are reported to remodel mitochondrial morphology and dynamics [74, 75]. Thus, integrating E2 and cellular functions, with PGC-1α, a coactivator of ERα via the LXXLL motif during JEV infection, addresses the important missing link [30, 33]. E2 and ERα signaling regulate PGC-1α through both transcriptional and post-translational mechanisms, linking estrogenic input to mitochondrial biogenesis [76]. It can be deduced that ERα can cooperate with PGC-1α to upregulate genes that increase energy demand during JEV replication, which may benefit replication. It has been shown that PGC-1α modulates the infection kinetics of the hepatitis C virus and the human rhinovirus [77–79]. Our data further suggested that inhibition of PGC-1α in neurons leads to downregulation of JEV, indicating the involvement of the ERα-PGC-1α axis during replication [80]. It is crucial for the inhibition of angiogenesis and promoting apoptosis.

While the N2a cell studies demonstrate that estrogen enhances JEV replication through the ERα-PGC-1α axis, this cellular pro-viral effect must be interpreted within the broader organismal context. Splenic protein analyses extended the brain findings to the peripheral immune compartment, indicating that estrogen not only aggravates central inflammation but also shapes the antiviral immune response. Histopathology and immunostaining data supported that intracerebral inoculation produced the expected moderate neuroinflammatory lesions.

This study reveals a sophisticated regulatory network in which estrogen signaling establishes a metabolic environment that enhances JEV replication at the cellular level. Sex hormones, including estrogens, are reported to be adversely affected by oxidative stress in neuronal cells [36]. The ERα-PGC-1α axis, which governs cellular homeostasis, provides a molecular framework for understanding differences in viral encephalitis outcomes and highlights the therapeutic potential of targeting hormone-metabolic cross-talk. The study could pave the way for selective ER modulators as therapeutic strategies for controlling JEV and other neurotropic viruses.

## Ethical statement

All the animal experiments were approved by the Institutional Animal Care and Use Committee of Assam Agricultural University, India. The study in the laboratory is conducted in accordance with the guidelines of the institutional biosafety committee, vide sanction IBSC/2021/MARCH/SK/BSBE/10.

## CRediT authorship contribution statement

**Kiran Singh:** Conceptualization, Data curation, Formal analysis, Investigation, Methodology, Visualization, Writing, original draft. **Vishnu Kumar (Computational work):** Conceptualization, Data curation, Formal analysis, Investigation, Methodology, Visualization, Writing - original draft. **Marek Bartkuhn (Computational work):** Supervision, Writing, review, and editing. **Sachin Kumar**: Resources, Supervision, Writing, review, and editing.

## Acknowledgment

Laboratory-based viral research is supported by the Indian Council of Medical Research (ICMR) (IIRP-2023-2444). Dr. Jumi Dutta and Dr. Archana Hazarika at Assam Agriculture University, Assam, support the animal experiment.

## Data Availability

The data sets generated during the study are provided within the paper and are available from the corresponding author upon appropriate request.

## References

[1] Kumar R, Kumar P, Singh MK, Agarwal D, Jamir B, Khare S, et al. Epidemiological Profile of Acute Viral Encephalitis. Indian J Pediatr. 2018;85:358–63.

[2] Kyaw AK, Ngwe Tun MM, Nabeshima T, Buerano CC, Ando T, Inoue S, et al. Japanese Encephalitis- and Dengue-Associated Acute Encephalitis Syndrome Cases in Myanmar. Am J Trop Med Hyg. 2019;100:643–6.

[3] Nath SR, Dasgupta A, Dubey D, Kokesh E, Beecher G, Fadra N, et al. Unraveling calcium dysregulation and autoimmunity in immune mediated rippling muscle disease. Acta Neuropathol Commun. 2025;13:11.

[4] Kumar R. Understanding and managing acute encephalitis. F1000Res. 2020;9.

[5] Gupta A, Bohara VS, Chauhan AS, Mohapatra A, Kaur H, Sharma A, et al. Alpha-synuclein expression in neurons modulates Japanese encephalitis virus infection. J Virol. 2024;98:e0041824.

[6] Gupta A, Bohara VS, Siddegowda YB, Chaudhary N, Kumar S. Alpha-synuclein and RNA viruses: Exploring the neuronal nexus. Virology. 2024;597:110141.

[7] Sharma KB, Vrati S, Kalia M. Pathobiology of Japanese encephalitis virus infection. Mol Aspects Med. 2021;81:100994.

[8] Liu S, Wang J, Yang J, Wen Y. The underlying mechanism of Guillain-Barre syndrome in a young patient suffered from Japanese encephalitis virus infection: a case report. Virol J. 2022;19:139.

[9] Muccilli SG, Schwarz B, Shue B, Jessop F, Shannon JG, Larson CL, et al. Mitochondrial hyperactivity and reactive oxygen species drive innate immunity to the yellow fever virus-17D live-attenuated vaccine. PLoS Pathog. 2025;21:e1012561.

[10] Liu X, Yan Q, Liu X, Wei W, Zou L, Zhao F, et al. PKM2 induces mitophagy through the AMPK-mTOR pathway promoting CSFV proliferation. J Virol. 2024;98:e0175123.

[11] El-Bacha T, Midlej V, Pereira da Silva AP, Silva da Costa L, Benchimol M, Galina A, et al. Mitochondrial and bioenergetic dysfunction in human hepatic cells infected with dengue 2 virus. Biochim Biophys Acta. 2007;1772:1158–66.

[12] Butler M, Chotiwan N, Brewster CD, DiLisio JE, Ackart DF, Podell BK, et al. Cyclin-Dependent Kinases 8 and 19 Regulate Host Cell Metabolism during Dengue Virus Serotype 2 Infection. Viruses. 2020;12.

[13] Dias SSG, Cunha-Fernandes T, Souza-Moreira L, Soares VC, Lima GB, Azevedo-Quintanilha IG, et al. Metabolic reprogramming and lipid droplets are involved in Zika virus replication in neural cells. J Neuroinflammation. 2023;20:61.

[14] Duarte G, Moron AF, Timerman A, Fernandes CE, Mariani Neto C, Almeida Filho GL, et al. Zika Virus Infection in Pregnant Women and Microcephaly. Rev Bras Ginecol Obstet. 2017;39:235–48.

[15] Wishart CL, Spiteri AG, Tan J, Macia L, King NJC. Metabolic Programming Drives Protective and Inflammatory Monocyte Fates in Viral Encephalitis. Adv Sci (Weinh). 2025;12:e05844.

[16] Nazneen F, Neupane B, Chen Y, Karim SU, You Z, Cui W, et al. Interleukin-17A signaling promotes CD8+ T cell cytotoxicity against West Nile virus infection through enhancing PI3K-mTOR-mediated metabolism. PLoS Pathog. 2025;21:e1013218.

[17] Zhao Y, Sui L, Pan M, Jin F, Huang Y, Fang S, et al. The segmented flavivirus Alongshan virus reduces mitochondrial mass by degrading STAT2 to suppress the innate immune response. J Virol. 2025;99:e0130124.

[18] Yang TC, Shiu SL, Chuang PH, Lin YJ, Wan L, Lan YC, et al. Japanese encephalitis virus NS2B-NS3 protease induces caspase 3 activation and mitochondria-mediated apoptosis in human medulloblastoma cells. Virus Res. 2009;143:77–85.

[19] Singh G, Singh K, Sinha RA, Singh A, Khushi, Kumar A. Japanese encephalitis virus infection causes reactive oxygen species-mediated skeletal muscle damage. Eur J Neurosci. 2024;60:4843–60.

[20] Agarwal A, Alam MF, Basu B, Pattanayak S, Asthana S, Syed GH, et al. Japanese Encephalitis Virus NS4A Protein Interacts with PTEN-Induced Kinase 1 (PINK1) and Promotes Mitophagy in Infected Cells. Microbiol Spectr. 2022;10:e0083022.

[21] Li M, Yang J, Ye C, Bian P, Yang X, Zhang H, et al. Correction to: Integrated Metabolomics and Transcriptomics Analyses Reveal Metabolic Landscape in Neuronal Cells During JEV Infection. Virol Sin. 2021;36:1693–4.

[22] Bohara VS, Deshmukh A, Kumar S. Pyruvate kinase M2 modulates Japanese encephalitis virus replication in neuronal cells. J Gen Virol. 2025;106.

[23] Bohara VS, Kumar S. Japanese encephalitis virus promotes its replication in neuronal cells by enhancing glycolysis via hypoxia-inducible factor-1alpha. Virology. 2026;614:110736.

[24] Tripathi S, Sengar S, Shree B, Mohapatra S, Basu A, Sharma V. An RBM10 and NF-kappaB interacting host lncRNA promotes JEV replication and neuronal cell death. J Virol. 2023;97:e0118323.

[25] Hsu JY, Hung CC, Tsou TP, Chen WC. Epidemiology and risk factors of Japanese encephalitis in Taiwan, 2010-2022. PLoS Negl Trop Dis. 2023;17:e0011421.

[26] Li X, Cui S, Gao X, Wang H, Song M, Li M, et al. The Spatio-temporal Distribution of Japanese Encephalitis Cases in Different Age Groups in Mainland China, 2004 - 2014. PLoS Negl Trop Dis. 2016;10:e0004611.

[27] Foster TC. Role of estrogen receptor alpha and beta expression and signaling on cognitive function during aging. Hippocampus. 2012;22:656–69.

[28] Kato S. Estrogen receptor-mediated cross-talk with growth factor signaling pathways. Breast Cancer. 2001;8:3–9.

[29] Manavathi B, Samanthapudi VS, Gajulapalli VN. Estrogen receptor coregulators and pioneer factors: the orchestrators of mammary gland cell fate and development. Front Cell Dev Biol. 2014;2:34.

[30] Guajardo-Correa E, Silva-Aguero JF, Calle X, Chiong M, Henriquez M, Garcia-Rivas G, et al. Estrogen signaling as a bridge between the nucleus and mitochondria in cardiovascular diseases. Front Cell Dev Biol. 2022;10:968373.

[31] Arnold S, Victor MB, Beyer C. Estrogen and the regulation of mitochondrial structure and function in the brain. J Steroid Biochem Mol Biol. 2012;131:2–9.

[32] Yang H, Yang R, Liu H, Ren Z, Kong F, Li D, et al. Synergism between PGC-1alpha and estrogen in the survival of endometrial cancer cells via the mitochondrial pathway. Onco Targets Ther. 2016;9:3963–73.

[33] Cao L, Li Y, Smirnov A, Voshtani R, Wang T, Shao C, et al. PGC-1alpha: key regulator of mitochondrial biogenesis and cellular differentiation in metabolic and regenerative tissues. Cell Biosci. 2025;16:9.

[34] Tcherepanova I, Puigserver P, Norris JD, Spiegelman BM, McDonnell DP. Modulation of estrogen receptor-alpha transcriptional activity by the coactivator PGC-1. J Biol Chem. 2000;275:16302–8.

[35] Maioli S, Leander K, Nilsson P, Nalvarte I. Estrogen receptors and the aging brain. Essays Biochem. 2021;65:913–25.

[36] Duong P, Tenkorang MAA, Trieu J, McCuiston C, Rybalchenko N, Cunningham RL. Neuroprotective and neurotoxic outcomes of androgens and estrogens in an oxidative stress environment. Biol Sex Differ. 2020;11:12.

[37] Yu Z, Su G, Zhang L, Liu G, Zhou Y, Fang S, et al. Icaritin inhibits neuroinflammation in a rat cerebral ischemia model by regulating microglial polarization through the GPER-ERK-NF-kappaB signaling pathway. Mol Med. 2022;28:142.

[38] Wei X, Hong L, Liang H, Ren K, Man W, Zhao Y, et al. Estrogen and viral infection. Front Immunol. 2025;16:1556728.

[39] Deguchi K, Kamada M, Irahara M, Maegawa M, Yamamoto S, Ohmoto Y, et al. Postmenopausal changes in production of type 1 and type 2 cytokines and the effects of hormone replacement therapy. Menopause. 2001;8:266–73.

[40] Herzog SK, Fuqua SAW. ESR1 mutations and therapeutic resistance in metastatic breast cancer: progress and remaining challenges. Br J Cancer. 2022;126:174–86.

[41] McCarthy M, Raval AP. The peri-menopause in a woman’s life: a systemic inflammatory phase that enables later neurodegenerative disease. J Neuroinflammation. 2020;17:317.

[42] Rishabh, Rohilla M, Bansal S, Bansal N, Chauhan S, Sharma S, et al. Estrogen signalling and Alzheimer’s disease: Decoding molecular mechanisms for therapeutic breakthrough. Eur J Neurosci. 2024;60:3466–90.

[43] Cerri S, Mus L, Blandini F. Parkinson’s Disease in Women and Men: What’s the Difference? J Parkinsons Dis. 2019;9:501–15.

[44] Morale MC, Serra PA, L’Episcopo F, Tirolo C, Caniglia S, Testa N, et al. Estrogen, neuroinflammation and neuroprotection in Parkinson’s disease: glia dictates resistance versus vulnerability to neurodegeneration. Neuroscience. 2006;138:869–78.

[45] Rocca WA, Faubion SS. Estrogen and dementia. Maturitas. 2022;165:120–1.

[46] Ulitzky L, Lafer MM, KuKuruga MA, Silberstein E, Cehan N, Taylor DR. A New Signaling Pathway for HCV Inhibition by Estrogen: GPR30 Activation Leads to Cleavage of Occludin by MMP-9. PLoS One. 2016;11:e0145212.

[47] Chen Q, Gouilly J, Ferrat YJ, Espino A, Glaziou Q, Cartron G, et al. Metabolic reprogramming by Zika virus provokes inflammation in human placenta. Nat Commun. 2020;11:2967.

[48] Gong S, Hao X, Bi Y, Yang C, Wang W, Mickael HK, et al. Hepatitis E viral infection regulates estrogen signaling pathways: Inhibition of the cAMPK-PKA-CREB and PI3K-AKT-mTOR signaling pathways. J Med Virol. 2021;93:3769–78.

[49] Khan S, Lew I, Wu F, Fritts L, Fontaine KA, Tomar S, et al. Low expression of RNA sensors impacts Zika virus infection in the lower female reproductive tract. Nat Commun. 2019;10:4344.

[50] Sharma P, Parmar K, Datta A, Ghosh B, Sarmah D, Borah A, et al. Recent progress and challenges of acute encephalitis syndrome: an update on pathogenesis and interventional approaches. Brain Disorders. 2025;Volume 19:100267.

[51] Hsieh JT, St John AL. Japanese encephalitis virus and its mechanisms of neuroinvasion. PLoS Pathog. 2020;16:e1008260.

[52] Ye X, Wang P, Song M, Zhang J, Li X, Wang X. Neuroinflammation and blood-brain barrier disruption in Japanese encephalitis virus infection: mechanisms, therapeutic targets, and intervention strategies. Arch Pharm Res. 2026;49:197–222.

[53] Yu SP, Ong KC, Perera D, Wong KT. Neuronal transcriptomic responses to Japanese encephalitis virus infection with a special focus on chemokine CXCL11 and pattern recognition receptors RIG-1 and MDA5. Virology. 2019;527:107–15.

[54] Das S, Basu A. Japanese encephalitis virus infects neural progenitor cells and decreases their proliferation. J Neurochem. 2008;106:1624–36.

[55] Arnal JF, Lenfant F, Metivier R, Flouriot G, Henrion D, Adlanmerini M, et al. Membrane and Nuclear Estrogen Receptor Alpha Actions: From Tissue Specificity to Medical Implications. Physiol Rev. 2017;97:1045–87.

[56] Alvarado MG, Schwartz DA. Zika Virus Infection in Pregnancy, Microcephaly, and Maternal and Fetal Health: What We Think, What We Know, and What We Think We Know. Arch Pathol Lab Med. 2017;141:26–32.

[57] Auriti C, De Rose DU, Santisi A, Martini L, Piersigilli F, Bersani I, et al. Pregnancy and viral infections: Mechanisms of fetal damage, diagnosis and prevention of neonatal adverse outcomes from cytomegalovirus to SARS-CoV-2 and Zika virus. Biochim Biophys Acta Mol Basis Dis. 2021;1867:166198.

[58] Das B, Dobrowolski C, Luttge B, Valadkhan S, Chomont N, Johnston R, et al. Estrogen receptor-1 is a key regulator of HIV-1 latency that imparts gender-specific restrictions on the latent reservoir. Proc Natl Acad Sci U S A. 2018;115:E7795–E804.

[59] Robinson DP, Lorenzo ME, Jian W, Klein SL. Elevated 17beta-estradiol protects females from influenza A virus pathogenesis by suppressing inflammatory responses. PLoS Pathog. 2011;7:e1002149.

[60] Malyala A, Kelly MJ, Ronnekleiv OK. Estrogen modulation of hypothalamic neurons: activation of multiple signaling pathways and gene expression changes. Steroids. 2005;70:397–406.

[61] Almey A, Milner TA, Brake WG. Estrogen receptors observed at extranuclear neuronal sites and in glia in the nucleus accumbens core and shell of the female rat: Evidence for localization to catecholaminergic and GABAergic neurons. J Comp Neurol. 2022;530:2056–72.

[62] Faure MC, Corona R, Roomans C, Lenfant F, Foidart JM, Cornil CA. Role of Membrane Estrogen Receptor Alpha on the Positive Feedback of Estrogens on Kisspeptin and GnRH Neurons. eNeuro. 2024;11.

[63] Pemberton K, Rosato M, Dedert C, DeLeon C, Arnatt C, Xu F. Differential Effects of the G-Protein-Coupled Estrogen Receptor (GPER) on Rat Embryonic (E18) Hippocampal and Cortical Neurons. eNeuro. 2022;9.

[64] Tsai MC, Lin SH, Hidayah K, Lin CI. Equol Pretreatment Protection of SH-SY5Y Cells against Abeta (25-35)-Induced Cytotoxicity and Cell-Cycle Reentry via Sustaining Estrogen Receptor Alpha Expression. Nutrients. 2019;11.

[65] Bandala C, Cardenas-Rodriguez N, Reyes-Long S, Cortes-Algara A, Contreras-Garcia IJ, Cruz-Hernandez TR, et al. Estrogens as a Possible Therapeutic Strategy for the Management of Neuroinflammation and Neuroprotection in COVID-19. Curr Neuropharmacol. 2023;21:2110–25.

[66] Upadhayay S, Gupta R, Singh S, Mundkar M, Singh G, Kumar P. Involvement of the G-Protein-Coupled Estrogen Receptor-1 (GPER) Signaling Pathway in Neurodegenerative Disorders: A Review. Cell Mol Neurobiol. 2023;43:1833–47.

[67] Wang C, Zhang F, Jiang S, Siedlak SL, Shen L, Perry G, et al. Estrogen receptor-alpha is localized to neurofibrillary tangles in Alzheimer’s disease. Sci Rep. 2016;6:20352.

[68] Torres T, Adam N, Mhaouty-Kodja S, Naule L. Reproductive function and behaviors: an update on the role of neural estrogen receptors alpha and beta. Front Endocrinol (Lausanne). 2024;15:1408677.

[69] Quesada A, Lee BY, Micevych PE. PI3 kinase/Akt activation mediates estrogen and IGF-1 nigral DA neuronal neuroprotection against a unilateral rat model of Parkinson’s disease. Dev Neurobiol. 2008;68:632–44.

[70] Fuentes N, Silveyra P. Estrogen receptor signaling mechanisms. Adv Protein Chem Struct Biol. 2019;116:135–70.

[71] Wijayaratne AL, McDonnell DP. The human estrogen receptor-alpha is a ubiquitinated protein whose stability is affected differentially by agonists, antagonists, and selective estrogen receptor modulators. J Biol Chem. 2001;276:35684–92.

[72] Grady SF, Pinto AK, Hassert M, D’Angelo JA, Brien JD, Arnatt CK. Selective estrogen receptor modulator, tamoxifen, inhibits Zika virus infection. J Med Virol. 2021;93:6155–62.

[73] Srivastava S, Khanna N, Mathur A, Chaturvedi UC. Role of calcium in neutrophil activation by Japanese encephalitis virus-induced macrophage derived factor. Acta Virol. 1994;38:111–6.

[74] Lee JK, Shin OS. Zika virus modulates mitochondrial dynamics, mitophagy, and mitochondria-derived vesicles to facilitate viral replication in trophoblast cells. Front Immunol. 2023;14:1203645.

[75] Chatel-Chaix L, Cortese M, Romero-Brey I, Bender S, Neufeldt CJ, Fischl W, et al. Dengue Virus Perturbs Mitochondrial Morphodynamics to Dampen Innate Immune Responses. Cell Host Microbe. 2016;20:342–56.

[76] Klinge CM. Estrogenic control of mitochondrial function. Redox Biol. 2020;31:101435.

[77] Shlomai A, Rechtman MM, Burdelova EO, Zilberberg A, Hoffman S, Solar I, et al. The metabolic regulator PGC-1alpha links hepatitis C virus infection to hepatic insulin resistance. J Hepatol. 2012;57:867–73.

[78] Yao W, Cai H, Li X, Li T, Hu L, Peng T. Endoplasmic reticulum stress links hepatitis C virus RNA replication to wild-type PGC-1alpha/liver-specific PGC-1alpha upregulation. J Virol. 2014;88:8361–74.

[79] Michi AN, Yipp BG, Dufour A, Lopes F, Proud D. PGC-1alpha mediates a metabolic host defense response in human airway epithelium during rhinovirus infections. Nat Commun. 2021;12:3669.

[80] Gorska-Ponikowska M, Kuban-Jankowska A, Eisler SA, Perricone U, Lo Bosco G, Barone G, et al. 2-Methoxyestradiol Affects Mitochondrial Biogenesis Pathway and Succinate Dehydrogenase Complex Flavoprotein Subunit A in Osteosarcoma Cancer Cells. Cancer Genomics Proteomics. 2018;15:73–89.

